# Illuminating the microbiome’s dark matter: a functional genomic toolkit for the study of human gut Actinobacteria

**DOI:** 10.1101/304840

**Authors:** Jordan E Bisanz, Paola Soto-Perez, Kathy N Lam, Elizabeth N Bess, Henry J Haiser, Emma Allen-Vercoe, Vayu M Rekdal, Emily P Balskus, Peter J Turnbaugh

## Abstract

Despite the remarkable evolutionary and metabolic diversity found within the human microbiome, the vast majority of mechanistic studies focus on two phyla: the Bacteroidetes and the Proteobacteria. Generalizable tools for studying the other phyla are urgently needed in order to transition microbiome research from a descriptive to a mechanistic discipline. Here, we focus on the Coriobacteriia class within the Actinobacteria phylum, detected in the distal gut of 90% of adult individuals around the world, which have been associated with both chronic and infectious disease, and play a key role in the metabolism of pharmaceutical, dietary, and endogenous compounds. We established, sequenced, and annotated a strain collection spanning 14 genera, 8 decades, and 3 continents, with a focus on *Eggerthella lenta*. Genome-wide alignments revealed inconsistencies in the taxonomy of the Coriobacteriia for which amendments have been proposed. Re-sequencing of the *E. lenta* type strain from multiple culture collections and our laboratory stock allowed us to identify errors in the finished genome and to identify point mutations associated with antibiotic resistance. Analysis of 24 *E. lenta* genomes revealed an “open” pan-genome suggesting we still have not fully sampled the genetic and metabolic diversity within this bacterial species. Consistent with the requirement for arginine during *in vitro* growth, the core *E. lenta* genome included the arginine dihydrolase pathway. Surprisingly, glycolysis and the citric acid cycle was also conserved in *E. lenta* despite the lack of evidence for carbohydrate utilization. We identified a species-specific marker gene and validated a multiplexed quantitative PCR assay for simultaneous detection of *E. lenta* and specific genes of interest from stool samples. Finally, we demonstrated the utility of comparative genomics for linking variable genes to strain-specific phenotypes, including antibiotic resistance and drug metabolism. To facilitate the continued functional genomic analysis of the Coriobacteriia, we have deposited the full collection of strains in DSMZ and have written a general software tool (ElenMatchR) that can be readily applied to novel phenotypic traits of interest. Together, these tools provide a first step towards a molecular understanding of the many neglected but clinically-relevant members of the human gut microbiome.

## Introduction

The isolation and sequencing of genomes from the human microbiome has been a major component of initiatives including the Human Microbiome Project (HMP) and Genomic Encyclopedia of Bacteria and Archaea (GEBA). This has led to the generation of thousands of genomes for the scientific community. These projects have focused on the sequencing of representative strains from the gut, skin, oral cavity, and other body habitats (HMP) (1) and systematically filling in holes in the tree of life (GEBA) (2). However, by definition these large-scale surveys favor breadth over depth and individual species are often poorly sampled; 60% of species in the HMP collection are represented by a single genome (1).

Strains of the same bacterial species are usually defined as having shared phenotypic characteristics (3) and, in the genomic era, 95% average nucleotide identity (4). When combined with the great diversity in gene content between bacterial strains often rendered by horizontal gene transfer (HGT), some have even questioned whether the species concept of bacteria is appropriate (5). Even isolates of the same strain which should be genetically identical can have meaningful variation affecting host interactions as has been observed with the variable presence of mucin-binding pili in *Lactobacillus rhamnosus* GG isolates (6). These observations highlight the importance of sampling the genetic and phenotypic diversity within specific clades of bacteria as a complementary approach to broad surveys. This is especially important for the development of microbiome-targeted diagnostics and interventions wherein low resolution markers are often used to infer genetic and phenotypic traits of communities (7). Here, we demonstrate the utility of conducting an indepth analysis of a single group of neglected, but clinically-relevant bacteria in these collections for which genetic tools are largely lacking. We focus on the Coriobacteriia class and in particular the species *Eggerthella lenta* due to their broad relevance for human health and disease, but the tools and approaches we describe could be readily applied to other taxonomic groups of interest.

Coriobacteriia are one of the most prevalent bacterial classes in the distal gut of adults, yet its low abundance and difficulty in isolation has led to a lack of biological material and tools for its study. The type genus of *Coriobacterium* was first isolated from the gut of the red soldier bug (8). Since then, the remaining genera have been primarily isolated from mammals suggesting a strong tropism for life as an intestinal symbiont (9). These bacteria, in particular the genera *Eggerthella* and *Paraeggerthella*, are often considered opportunistic pathogens due to their common isolation from bacteremia patients(10–12). More recently, microbiome surveys have revealed positive associations between these bacteria and type 2 diabetes, cardiovascular disease, and rheumatoid arthritis though causative links remains to be established (13–15).

Members of the Coriobacteriia are also versatile in the metabolism of pharmaceutical, dietary, and endogenous compounds. *E. lenta*, a focus of this work, inactivates the cardiac drug digoxin (16–18) and metabolizes bile acids (19, 20). This class also plays a key role in the production of small molecules that are similar to estrogen in their chemical structure and function (referred to as phytoestrogens). *E. lenta* contributes to the multi-step biotransformation of plant-derived polyphenolic lignans to estrogen-like compounds (phytoestrogens) (21, 22), whereas *Slackia isoflavoniconvertens*, a close relative, metabolizes the plant-derived isoflavone daidzein to the phytoestrogen equol and 5-hydroxy-equol (23)

The primary goal of this work was to assemble a collection of isolates from the Coriobacteriia, and in particular the species *E. lenta*, to be sequenced and made publicly available to facilitate improved understanding of this understudied but clinically-relevant group of human gut bacteria. Through the use of these genomes, we have designed an assay for the sensitive and selective quantification of *E. lenta* and its effectors from fecal samples. In addition to releasing their genome sequences, a user-friendly tool, ElenMatchR has been developed to facilitate gene-trait matching experiments (jbisanz.shinyapps.io/elenmatchr/) which we validate by uncovering antibiotic resistance genes naturally occurring within the *E. lenta* species and with previously described genes for drug inactivation (16, 18).

## Results and Discussion

### Genome Sequencing

Through public collections (n=22), and novel isolation (n=28), we collected 50 isolates for study (**Supplemental Table 1**). This collection represents 14 constituent genera of the Coriobacteriia (9, 24) including 30 *Eggerthella lenta* isolates. While most isolates were derived from human-associated communities, a single isolate is derived from the murine cecum (*Parvibacter caecicola* DSM 22242). Strains range in year of isolation from 1938 to 2016 and represent 8 countries over 3 continents from regions of variable Coriobacteriia prevalence in human gut microbiomes ranging from 86.8 to 96.4% (**Supplemental Figure 1**). Of these 50 isolates, 33 were without a sequenced public genome leading to us to initiate a sequencing project over the course of multiple years using variable sequencing strategies as new strains became available. A total of 33 new high-quality draft genomes were generated (62 ±18 total scaffolds, 152 ± 62 kb N50, 8.6 ± 2.4 scaffolds L50 [mean ± sd]). More complete genome assembly statistics are available on a per-genome basis in **Supplemental Figures 2AB, and Supplemental Table 2**. Genomes were considered to be free of contaminants based on pre-assembly filtering sequencing reads for human, phiX174, and Illumina/NEBNext adapter sequences (see methods), and an approximate Gaussian distribution in GC content (**Supplemental Figure 2C**). In the process of analysis, an additional public genome (*Clostridium difficile* F501: PRJNA85919) was uncovered sharing significant nucleotide homologies to *E. lenta*. On closer inspection, it was determined on the basis of GC distribution and 16S rRNA content, that this genome is heavily contaminated by high GC content DNA consistent with an *E. lenta* strain (**Supplemental Figure 3**). Recovery of 16S rRNA sequences from contigs had 100% identity to *E. lenta* DSM 2243 and 99% identity to *Parabacteroides distasonis* ATCC 8503 suggesting an impure DNA preparation. The genome has been flagged as an anomalous assembly; however, an issue that is being repaired by Genbank staff resulted it in still appearing in BLAST databases. This contamination has led to inappropriate conclusions by other groups who would be justly unaware of this contamination (25).

### Taxonomic and phylogenetic description

The taxonomy of the Coriobacteriia was recently amended separating the constituent members into 3 families that were formerly all within the *Coriobacteriaceae* to include the additional families *Eggerthellaceae* and *Atopobiaceae* (24). This analysis reflects the current list of prokaryotic names with standing nomenclature (26). Given that many of the strains sequenced were not originally assigned a species (examples: DSM 11767 and DSM 11863) or were novel isolates, we sought to validate their taxonomy on the basis of genome-wide average nucleotide identity (ANI). ANI is an *in silico* alternative to traditional DNA-DNA hybridization (DDH) wherein 95% ANI is generally accepted as the species boundary corresponding to 70% DDH. All strains of *E. lenta* (taxonomy based on 16S rRNA sequence or culture collection identification) were confirmed with ANI varying from between 97.98 to 99.64% (98.65 ± 0.32 mean ± sd) within the species *E. lenta*, and 81.96 to 91.40% (84.83 ± 1.94) to isolates outside the species (**Figures 1AB**). ANI was also applied in combination with isolation metadata to dereplicate isolates removing suspect clonal isolates (>99.999% ANI, n=3 strains with clonal isolates, n=24 total dereplicated strains).

**Figure 1.**
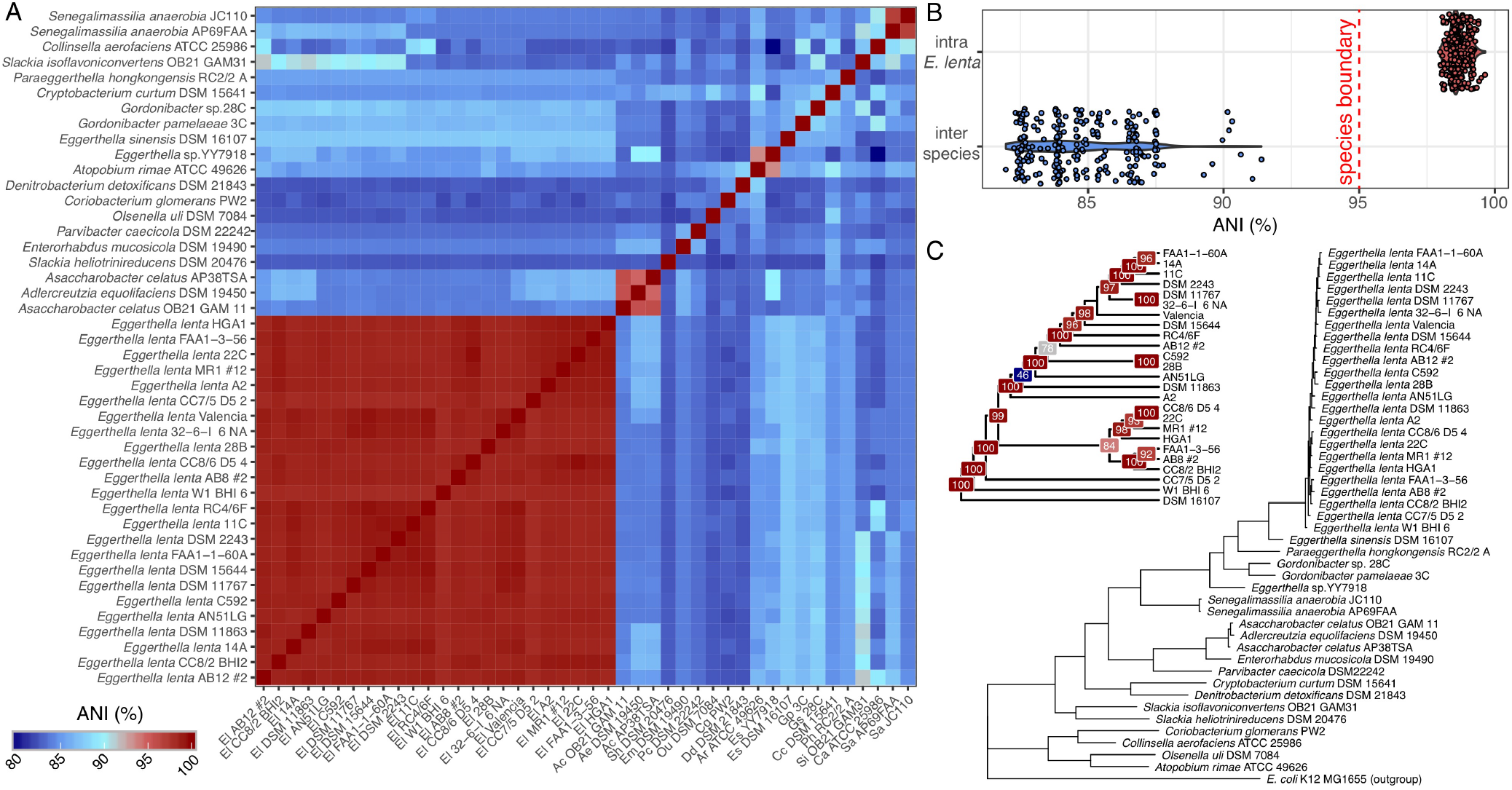
Genomic analysis supports a revised taxonomy for the Coriobacteriia. (**A**) Average nucleotide identity (ANI) was calculated between all pairs of dereplicated Coriobacteriia strains demonstrating clear speciation boundaries (**B**) between *Eggerthella lenta* (n=24) and other Coriobacteriia (n=20) validating taxonomic assignments to *E. lenta*. (**C**) Phylogenetic tree of the Coriobacteriia demonstrates the mis-assignment of *Eggerthella* sp. YY7918 to the genus *Eggerthella* and uncertainty in the boundaries between *Asaccharobacter* and *Adlercreutzia (E. coli* as out-group). The inset shows a cladogram of *E. lenta* with *E. sinensis* as out-group and bootstrap values at internal nodes.

This analysis highlights an issue relating to the assignment of the *Eggerthella* YY7918 strain to its genus given the lack of homology to *Eggerthella* genus (84.50 to 87.21%, 85.88 ± 1.01 mean ± sd). Its location in a phylogenetic tree also clearly positions it outside of the genus (**Figure 1C**) and its GC content is only 56% deviating far from the 64% and 65% of *E. lenta* and *E. sinensis*. Given these data, and the nearest match of 93% to *Gordonibacter* spp. by 16S rRNA sequence, it is proposed that *Eggerthella* YY7918 represents a novel genus for which it is the type species. The inclusion of this genome into databases for taxonomic assignment of short reads from microbiome studies, including the default database for MetaPhlAn2 (27), may lead inappropriate inferences of abundances of the *Eggerthella* genus.

We also report a close similarity between the genera *Asaccharobacter* and *Adlercreutzia* (**Figures 1AB**, **Supplemental Table 3**), two genera described in May 2008 (28, 29), and whose type species’ partial 16S rRNA sequences share 99% identity with 99% coverage (Genbank accessions: AB306661 and AB2666102). This suggests the existence of these two separate genera is inappropriate and that they should be collapsed to a single genus. Due to these discrepancies in the accepted nomenclature, strains AP38TSA and OB21GAM11 have been deposited with their original taxonomic identification of *Asaccharobacter celatus*.

### Identification of intra-isolate genetic polymorphisms

The type strain of *E. lenta* (DSM 2243 = ATCC 25559 = VPI 0255 = 1899 B) was first sequenced by Saunders *et al*. in 2009 (30) using a combination of Sanger and pyrosequencing as part of the GEBA project (2). Given that phenotypically relevant genetic mutations can occur between isolates of the same strain (6), we resequenced our laboratory strain of DSM 2243 (referred to herein as UCSF 2243), and fresh isolates from the ATCC and DSMZ culture collections, comparing them to the original contiguous reference assembly of Saunders *et al*. (30). All strains were sequenced at ultra-high depth and resulting reads were mapped against the reference assembly demonstrating that no elements have been lost from the genome (median per-base coverage of 391, range 33 to 5942; **Figure 2A**). Elevated coverage was observed in the region around a putative prophage (3.02-3.06 Mbp) suggesting an active prophage is present in all 3 stocks.

**Figure 2.**
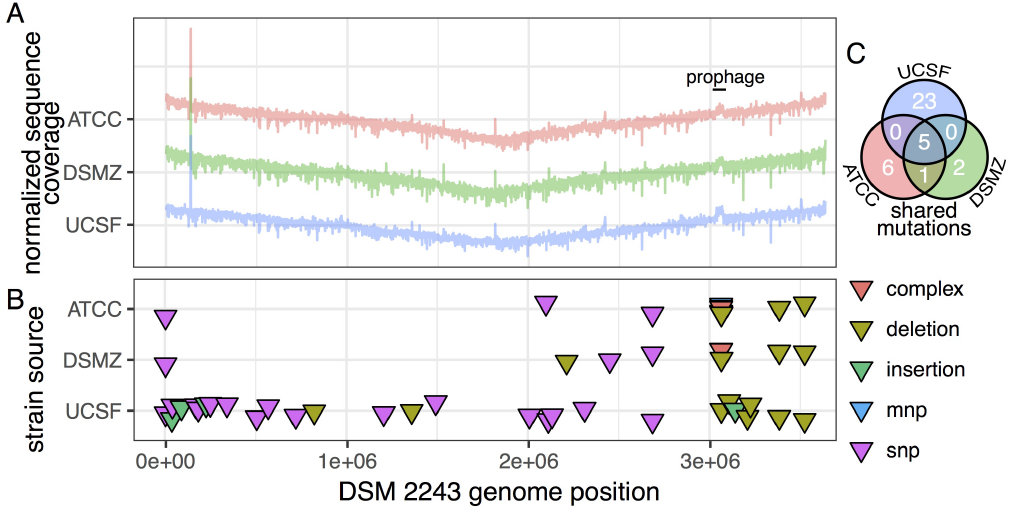
Identification of genetic polymorphisms in the *Eggerthella lenta* type strain. (**A**) Normalized sequence coverage indicates a minimum of 33-fold coverage (median: 391, range: 33-5964) across the entire contiguous reference genome assembly (CP001726) suggesting no loss of genomic islands. (**B**) Genetic variation between re-sequenced isolates suggests 5 variants (**C**) represent errors in the reference genome while the lab strain (UCSF) contains an additional 23 unique variants.

Next, the alignments were used to call variants between the newly sequenced strains and the reference sequence (**Figures 2BC**, **Supplemental Table 4**). A total of 5 polymorphisms were found in all sequenced isolates and it is likely that these represent errors in the reference sequence (**Figure 2C**). 3 of 5 shared polymorphisms are insertion/deletions in homopolymeric runs which is indicative of 454-type error while a single missense variant is found in the DNA polymerase III beta-subunit. Interestingly, the ATCC isolate contains a missense variant in a putative beta-lactamase (Elen_1788) resulting in a glycine to aspartic acid substitution outside of the predicted beta-lactamase domain (Pfam: PF00144). A literature review of antibiotic susceptibility analyses and treatment of *Eg-gerthella*-associated infections suggested that the beta-lactam antibiotics penicillin G and ampicillin have been used for treatment (Supplemental Table 5) leading us to examine the relevance of this SNP. We found that this variant in the ATCC 25559 strain was associated with a modest 2-4 fold increase in the minimum inhibitory concentrations of both penicillin G and ampicillin (**Supplemental Table 6**). According to the current clinical guidelines (31), this subtle shift would be sufficient to conclude that the ATCC 25559 isolate is resistant to penicillin G while the other two isolates are not. Further genetic studies are necessary to validate the causal role of this mutation in the antibiotic resistance phenotype, but these data emphasize the difficulties of relying on publicly available reference genomes for functional studies.

### Analysis of core and variable genes within *Eggerthella lenta*

To facilitate comparative genomic analysis, we analyzed gene content across the *E. lenta* strain collection by clustering all predicted coding sequences into orthologous groups (OGs) (32). Rarefaction analysis suggests an open (unsaturated) pan-genome (**Figure 3A**) with >5,871 OGs and a core of 1,832 OGs. At 24 dereplicated *E. lenta* isolate genomes, 74 new OGs (range:16-178) were discovered with each additional genome sequenced. To reconstruct core and variable metabolic pathways, we mapped the pan-genome to curated KEGG orthologous groups (KOs) with 44.3% of genes across the collection being assigned to an orthology (**Supplemental Figure 4**).

**Figure 3.**
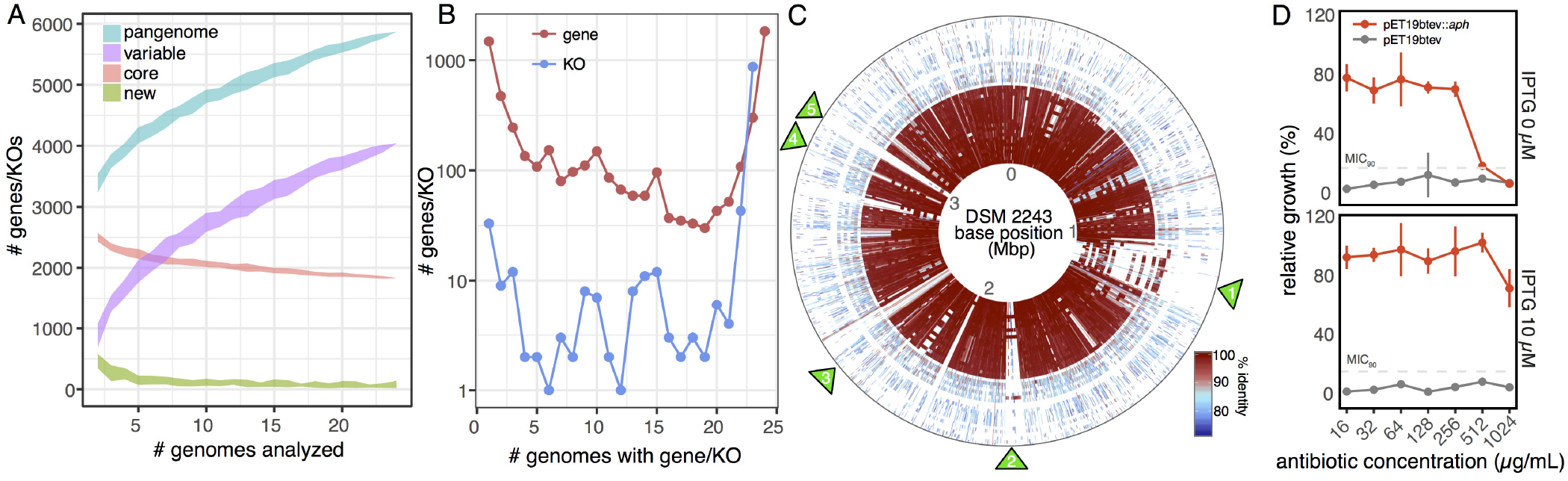
*Eggerthella lenta* has a large and unsaturated pan-genome. (**A**) Analysis of OGs across 24 strains of *E. lenta* demonstrates an open pan-genome (still increasing in size) with a current core genome of 1,832 OGs representing 31.2% of the pan-genome. A median of 71 new OGs (range 17-178) are discovered with every additional genome. Lines denote mean±sd. (**B**) OG and KEGG orthologous group (KO) conservation among *E. lenta* strains. A large proportion of genes and functional annotations (KOs) display inter-strain variability (69% and 16% respectively observed variably across genomes). (**C**) BLAST comparison of all strains against the *E. lenta* DSM 2243 reference genome. Each track represents the genome of a comparator strain (ordered by ANI) compared against the DSM 2243 reference genome demonstrating regions of homology across all strains. Green triangles denote regions of interest (ROIs, see main text). (**D**) Expression of a putative kanamycin resistance protein (Aph) in *E. coli* Rosetta reveals increased MIC compared to vector controls under leaky basal expression of the T7-lac promoter (0 μM IPTG) and increased resistance with low-level induction (10 μM IPTG).

Metabolic reconstruction of the *E. lenta* core genome questions the proposed reliance of this bacterium on amino acid catabolism for growth. As expected, the arginine dihydrolase (or arginine deiminase) pathway was present in all strains which explains the need for arginine supplementation to promote in vitro growth (16). However, the entire backbone of glycolysis was conserved along with all but 1 enzyme of the citric acid cycle (**Supplemental Figure 4**). Further more, genes including GAPDH are amongst the most highly expressed genes during exponential growth as measured by RNA-seq (16). These results conflict with growth assays suggesting that *E. lenta* is asaccharolytic (30). Metabolomic and isotope tracking studies are necessary to more definitively test the activity of these enzymes during *in vitro* or *in vivo* growth, and the relative contributions of carbohydrate versus amino acid metabolism to cellular metabolism.

Another notable pathway within the *E. lenta* core genome is predicted to convert isoprenoids into the carotenoid lycopene (**Supplemental Figure 4**). This pathway could provide the molecular basis for the production of a red-brown pigment in *E. lenta* colonies (**Supplemental Figure 5**). In practice, the formation of the pigment is highly variable in the laboratory; however, if the factors that regulate carotenoid production could be known, this feature may help in the more rapid isolation of additional strains as no selective media are currently available (9). Furthermore, bacterial biosynthesis of carotenoids in the gastrointestinal tract may have important downstream consequence for host pathophysiology (33).

Analysis of the variable component of the *E. lenta* genome highlighted multiple genomic islands indicative of extensive horizontal gene transfer (HGT) within this bacterial lineage. Genome-wide mapping of each strain against the type strain revealed multiple large-scale gene gain/loss events (Figure 3C). Two regions of interest (ROI1 and ROI2, Figure 3C) reflect potential hot spots of HGT. The 150 Kbp ROI1 contains a type IV secretion system (1.08 Mbp) implicated in HGT and virulence (34) in addition to a TN916 transposase, recombinase, and phiRv2 integrase. The region has partial homology to many other strains of *E. lenta* but also to *S. isoflavoniconvertens, Gordonibacter* spp., and *Senegalemas-silia anaerobia* indicative of both intra-species and intra-class HGT. The 54 Kbp ROI2 has been previously described due its association with an 8 bp GAGTGGGA motif recognized by P4 integrases present within a host GMP synthase gene (35). It is associated with genes involved in transposase activity and metal-resistance. ROI3 contains a strain-variable Type I-C CRISPR-cas machinery. ROI4 is home to the digoxin-metabolizing *cgr* operon (16, 18) and ROI5 contains a putative *E. lenta* prophage.

As an initial step towards developing selective markers for *E. lenta* genetic manipulation, we searched for antibiotic resistance genes found in only a single strain. We identified a gene with perfect sequence similarity to kanamycin kinase (aminoglycoside phosphotransferase, antibiotic resistance ontology 300264). This gene (referred to herein as *E. lenta aph*) was unique to *E. lenta* DSM 11767, which had significantly increased kanamycin resistance relative to the rest of the strain collection (**Supplemental Table 5**). Heterologous expression of *E. lenta aph* in *E. coli* resulted in an induction dose-dependent increase in antibiotic resistance (**Figure 3D**). Kanamycin resistance is widely used as a marker for bacterial genetics, making this gene a valuable addition to our growing genetic toolkit for *E. lenta*.

### Development of a duplexed assay for quantifying *E. lenta* species and gene abundance

Given the absence of a selective and/or differential culture media for *E. lenta* and the expensive, often low resolution, and time-consuming nature of taxonomic identification by 16S rRNA gene and shotgun metagenomic sequencing, we sought to develop a highly sensitive qPCR-based assay for *E. lenta*. This assay should simultaneously quantify *E. lenta* and the *cgr2* gene, which is responsible for the inactivation of the cardiac drug digoxin (18), with the ability to be combined with other effector-specific probes. In this way, we could develop a rapid, affordable, and sensitive assay that could be used to estimate *E. lenta* abundance and help to estimate the drug-metabolizing capacity of each individual’s microbiome.

To accomplish these aims, a multiplexed double-dye probe assay was designed. We used our comparative genomics approach to identify single-copy chromosomal genes that could serve as surrogate markers of the *E. lenta* strains by screening for highly conserved genes present in *E. lenta* but absent in the remainder of the Coriobacteriia and reporting no hits in the NCBI non-redundant database to other species. This led to the identification of a gene we termed *elnmrk1*, a luxR family transcriptional regulator (Elen_1291, ACV55260.1). Using a curated collection of processed shotgun metagenomes (36), we analyzed the correlation between *elnmrk1* and *E. lenta* mOTU (metagenomic operational taxonomic unit) abundance (37). We found a strong correlation (r=0.97 Pearson correlation, **Figure 4A**) validating that it is an appropriate surrogate marker for quantification. There is also not strong evidence of an *elnmrk1* negative *E. lenta* population. qPCR primer and probes (**Supplemental Table 7**) were designed to both *elnmrk1* and *cgr2* and were validated against a dilution series of a pure culture of *E. lenta* DSM 2243 (**Figure 4B**) as well as deeply-sequenced human metagenomes that had previously been taxonomically analyzed (**Figure 4B inset**). While the DNA extraction method will impact recovery and detection, the demonstrated method using mechanical lysis and magnetic bead recovery yielded linear recovery of pure cultures down to nearly 100 CFU/mL. Even in the presence of a second more highly abundant and easily lysed organism (*E. coli* K12 MG1655), sensitive quantification was possible without specificity issues commonly associated with 16S rDNA targeting assays.

**Figure 4.**
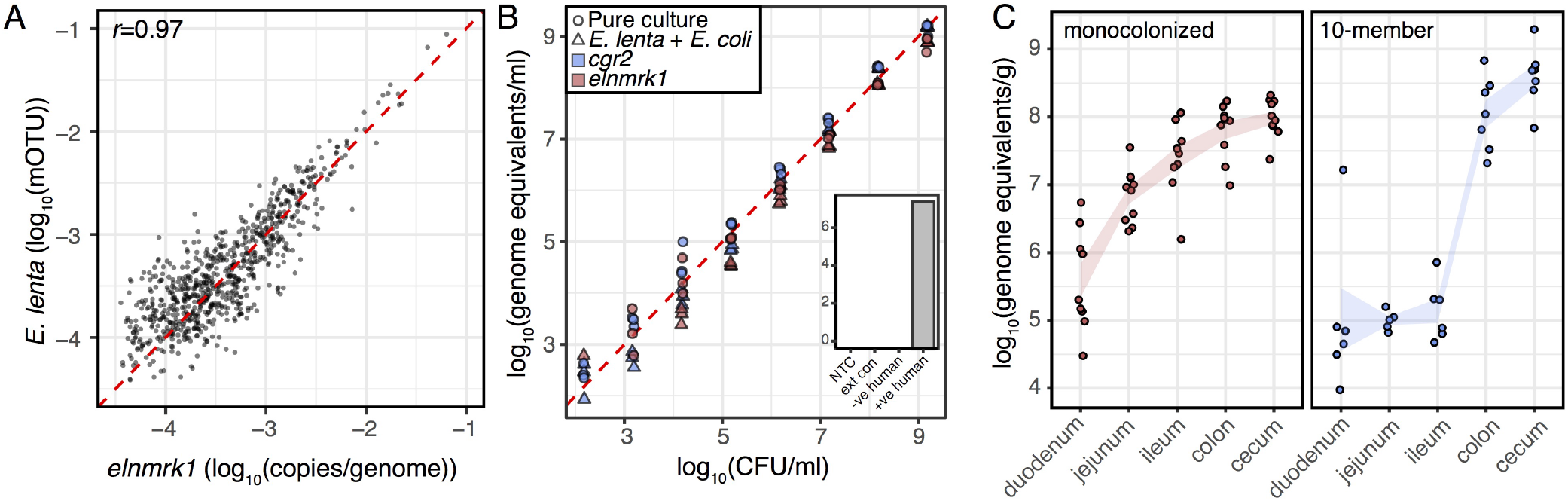
Validation of a duplexed qPCR assay for quantifying *Eggerthella lenta* and specific genes of interest. (**A**) Correlation of *elnmrk1* and *E. lenta* abundances in metagenomes reveals a high degree of correlation (p<0.001, r=0.97) and little evidence of an *elnmrk1*-negative *E. lenta* population in 760 human GI tracts (red line denotes expected linear relationship). (**B**) Validation of assay sensitivity demonstrates a practical quantification limit of 1400 genome equivalents and that the presence of a second more abundant, and easily lysed, organism (4E9 CFU/mL *E. coli* K12 MG1655) does not prevent the recovery and quantification of *E. lenta* (inset controls: no-template control (NTC), blank DNA extraction control (ext con), and negative and positive human controls [based on shotgun metagenomic sequencing, see methods]). (**C**) Quantification of *E. lenta* DSM 2243 in gnotobiotic mice mono-colonized or colonized with a 10-member synthetic community (**Supplemental Table 7**). *E. lenta* is detectable along the entire length of the gastrointestinal tract with highest levels observed in the colon and cecum. Points represent individual animals (n=6-10/group) and lines represent mean±SEM.

We next used this assay to quantify *E. lenta* DSM 2243 along the GI tract of both mono-colonized and 10-member mock-community colonized Swiss Webster mice. This analysis demonstrates that *E. lenta* can colonize along the entire murine GI tract, albeit at very low abundances (**Figure 4C**). The composition of the mock community (as determined from V4 16S rRNA gene amplicon sequencing and total 16S rRNA qPCR) is given in **Supplemental Table 8** providing corroboration of the qPCR results. These data and **Figure 4A** indicate that *E. lenta* is found predominantly at very low abundance in both humans and mice, which may explain why its prevalence has been so underestimated in the microbiome. Both qPCR- and metagenome-based prevalence estimates indicate that the *E. lenta* is a prevalent member of the human gut microbiome: 74.7% and 41.5%, respectively (18). Given the clear spectrum of colonization by *E. lenta*, we are actively pursuing the genetic and environmental determinants which govern colonization by this species.

### Gene-phenotype matching in *E. lenta* via comparative genomics

A major motivation for assembling this collection of strains and genomes was to facilitate the discovery of new effectors of metabolism of pharmaceutical, dietary, and endogenous compounds. To make this resource available to the scientific community, the approach and database from which it draws have been made publicly available via a tool we describe as ElenMatchR (jbisanz.shinyapps.io/ElenMatchR). Drawing on the logic of other tools such as PhenoLink (38), gene presence/absence is binarized/dereplicated, and used as the input predictor variable for a random forest classifier against user-provided phenotypes or traits (**Supplemental Figure 7**). Random forests are advantageous when data is particularly noisy and/or gene networks or multiple effectors are present. At the current time only categorical phenotypes are accepted; however, regression will be supported for continuous phenotypes in the future. The predictive importance (mean decrease in gain in node impurity), is used to extract and rank features. The results of the classifier are returned in addition to: (i) a scatter plot of feature accuracy, (ii) a heat map of gene presence and absence, (iii) candidate nucleotide and amino acid sequences, and (iv) a visualization of the phenotype on a pan-genome and phylogenetic tree.

For a test dataset, since we were aware of pre-existing variation in antibiotic resistance patterns to kanamycin, we opted to assay well-described phenotypes and effectors of antibiotic resistance using both broth microdilution minimal inhibitory concentration assays and Epsilometer (Etest) strips. Antibiotics were selected based on a literature review of susceptibility testing and clinically relevant antibiotics (**Supplemental Table 5**) leading to the screening of 8 antibiotics that are useful for genetic engineering and/or clinically use including the previously mentioned beta-lactams and kanamycin (**Supplemental Table 6**). Tetracycline resistance provided a binary phenotype by broth microdilution (**Figure 5A**) with 10 strains being resistant (MIC>32 μg/mL) and 9 strains being sensitive (MIC=4 μg/mL). This was expanded to 14 resistant strains and 12 sensitive strains via Etests (**Figure 5B**). Using clustering thresholds of 60% amino acid identity and a minimum 80% query coverage for generating orthologous gene clusters (OGs), and after removal of uninformative OGs (present or absent in all strains assayed) a total of 8,853 OGs were considered. After collapsing co-occuring OGs, 1,150 features were used as the input for the classifier. On inspecting variable importance, it is clear that a single OG (3652) provides the most discriminative value between classes (**Figure 5C**). Visualization of presence/absence shows that it is present in all 14 resistant strains, but absent in all sensitive strains (**Figure 5D**). Further analysis of this orthologous cluster of genes, annotated as tetracycline resistance protein TetO, shows that it is comprised of two subclusters with 67.3% amino acid identity to each other (**Figure 5E**). All 14 tetracycline resistance proteins were then annotated against the Comprehensive Antibiotic Resistance Database revealing the two subclusters to represent the related *tetO* and *tetW* ribosomal protection proteins with an average blast identity of 99.4% (range 98.28 to 99.7, ARO 3000190 and 3000194 respectively) (39). To validate this gene, a putative *tetW* from *E. lenta* DSM 11767 was cloned under the T7-lac promoter of pET19btev to be heterologously expressed in *E. coli* Rosetta. Expression of *tetW* conveyed resistance to tetracycline (**Figure 5F**) increasing the MICs from 1 μg/mL to ≥ 16 μg/mL in an induction-dose dependent manner. These genes, together with the kanamycin resistance gene identified in our pan-genome analysis, represent suitable antibiotic selection markers for the purposes of establishing tools for the genetic manipulation of *E. lenta*.

**Figure 5.**
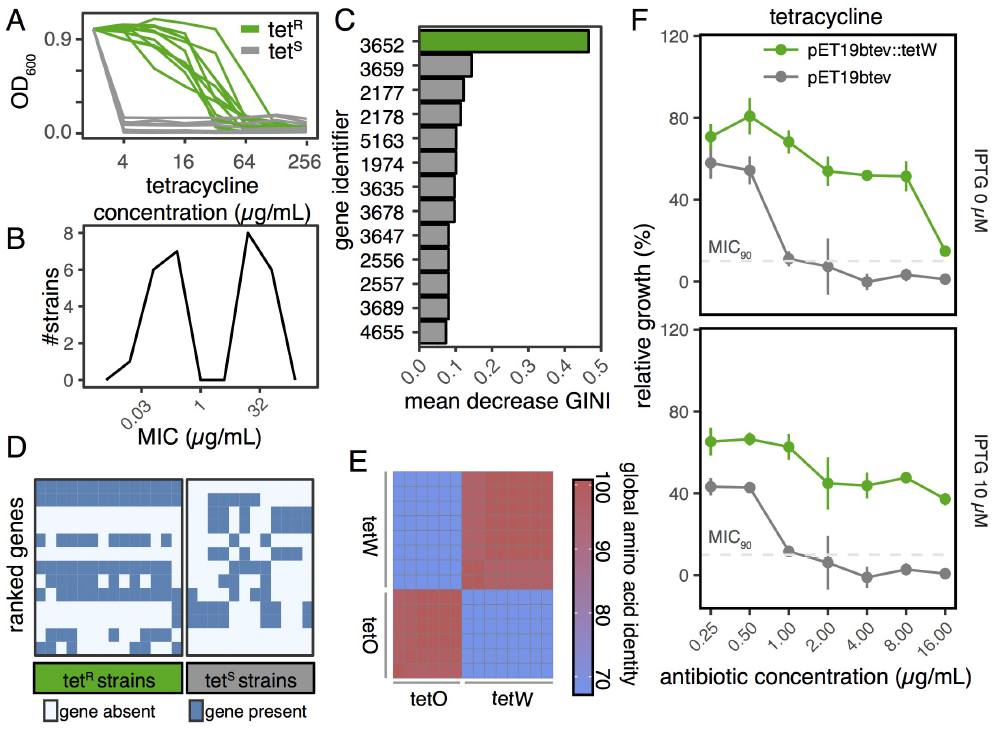
Comparative genomics enables the discovery of antibiotic resistance genes. (**A**) Broth microdilution demonstrated a binary tetracycline resistance phenotype with 10 resistant (green lines) and 9 sensitive (grey lines) strains. (**B**) Epsilometer (Etest) analysis of minimal inhibitory concentration (MIC) on solid media increases counts to 14 resistant and 12 sensitive strains with a bimodal distribution. (C) Gene importance scores (mean decrease gain in node impurity as determined by random forest, top 10 shown) identifies a single orthologous group (3652) with elevated importance (4-fold higher than next highest ranked OG). (**D**) Gene presence/absence (heat map) shows that gene group 3652 is present in all resistant strains while completely absent in sensitive strains. Annotation of the cluster identifies its members as *tetO/tetW* ribosomal protection proteins. (E) Clustering of tetracycline resistance proteins in cluster 3652 reveals two subfamilies of TetO and TetW proteins. (F) Expression of *tetW* results in an increased MIC compared to vector controls under leaky basal expression of the T7-lac promoter (0 μM IPTG) and increased resistance with low-level induction (10 μM IPTG).

Finally, we showed that ElenMatchR can also be readily applied to identify genes for drug metabolism as it correctly identifies the cardiac glycoside reductase (*cgr*) operon responsible for digoxin reduction (16) as well as an expanded loci with which it co-occurs termed the cgr-associated gene cluster (18) (**Supplemental Figure 8**). Both tetracycline resistance and digoxin metabolism datasets are included with ElenMatchR for demonstration purposes.

## Conclusions

Mechanistic studies into the basic biology of the Coriobacteriia coupled to translational studies in preclinical animal models and human cohorts are critical to gaining a greater understanding of the role of these intestinal symbionts in human health and disease. The functional genomic tools described herein have been developed to be shared with the greater scientific community. This includes our open source software package ElenMatchR (jbisanz.shinyapps.io/elenmatchr/), the deposition of these genome assemblies to Genbank, and the availability of novel isolates from the German Collection of Microorganisms and Cell Cultures (DSMZ).

Our in-depth analysis of the *E. lenta* pan-genome suggest that we are just beginning to scratch the surface of the genetic and biochemical potential of this species with many more variable genes remaining to be discovered. Despite our partial view of the full microbial genetic and metabolic diversity within this lineage, we have demonstrated the utility of genome mining, comparative genomics, and nucleotide-resolution analyses in elucidating the genetic basis of antibiotic resistance and drug metabolism. The identified antibiotic resistance genes provide a valuable first step towards developing tools for the genetic manipulation of Coriobacteriia. By making both the tool, and its pre-computed underlying databases available, we are openly facilitating others to work in this field and to modify the tool as they see fit. Ongoing studies in our lab are focused on using this framework to provide mechanistic insights into the role of *E. lenta* and other Coriobacteriia in metabolism, immunity, microbial interactions, and host pathways relevant to disease.

Translational studies in human cohorts will be facilitated by the development of a sensitive assay for the quantification of *E. lenta* and genes of interest from stool samples. This assay is a marked improvement from our earlier protocol (16) due to increased sensitivity and specificity as well as the ability to distinguish *E. lenta* strains within a single reaction. Additional studies are warranted to test the proof-of-principle for using this approach as a rapid diagnostic test to predict drug response. As new genetic effectors are discovered for other host or microbial phenotypes of interest, they could be multiplexed into the current assay for multiple marker detection. Together, these results highlight that moving beyond shallow coverage of gut bacterial taxonomic groups will improve our collective understanding of the role of the microbiome in health and disease. Our data emphasizes that single type strains and their genomes do not represent the broader scope of genotypic and phenotypic diversity within a given clade. In some cases, even single nucleotide polymorphisms appear to have phenotypic consequences, necessitating genome-wide nucleotide resolution and routine re-sequencing of culture collections. Progress towards microbiome-based precision medicine will likely require this level of specificity for accurate diagnostic and prognostic markers. More broadly, this work represents a generalizable template for the dissection of additional neglected taxonomic groups with the longterm goal of understanding the molecular mechanisms responsible for changes in the structure and function of host-associated microbial communities.

## Acknowledgements

We are indebted the UCSF Gnotobiotic Core Facility; Arianne Panzer and other members of the Lynch lab (UCSF); Michelle Daigneault for technical assistance (UoG), and An Luong, Niki Arab, and the other members of the Turnbaugh lab for technical assistance. The Turnbaugh lab is supported by the National Institutes of Health (R01HL122593). PJT is a Chan Zuckerberg Biohub investigator and a Nadia’s Gift Foundation Innovator supported, in part, by the Damon Runyon Cancer Research Foundation (DRR-42-16), the UCSF Program for Breakthrough Biomedical Research (partially funded by the Sandler Foundation), and the Searle Scholars Program. Isolation of new strains was partly funded by a Natural Sciences and Engineering Research Council of Canada (NSERC) Discovery Grant to EAV. JEB was supported by a NSERC postdoctoral fellowship. KNL is supported by the Canadian Institutes of Health Research Fellowship program.

## Author Contributions

PJT, HJH, EAV, VMR, and EPB were involved in the isolation of bacterial strains. JEB, KNL, HJH, PJT, and VMR were responsible for the library preparation and sequencing of genomes. JEB assembled, annotated, and analyzed genomes, deposited the genomes, and wrote ElenMatchR. JEB and PJT wrote the manuscript with input from all co-authors.

## Methods

### Strain Isolation and Culture

Novel strains were isolated in multiple medias. Briefly, this includes Eubacterial minimal media supplemented with 1% w/v arginine, 50 μg/mL hygromycin, and 25 μg/mL nalidixic acid; brain heart infusion agar (BHI) supplemented with 1% arginine; fastidious anaerobe agar (FAA); and nutrient agar. All strains were isolated under anaerobic conditions at 37°C. Routine culturing was carried out under anaerobic conditions (Coy Lab Products) using the appropriate media listed in **Supplemental Table 1**. Media used for routine experimentation were BHI+ (BHI with 1% arginine), BHI++ (BHI with 1% arginine, 0.05% L-cysteine-HCl, 1 μg/mL vitamin K, 5 μg/mL hemin, and 0.0001% w/v resazurin), or TSA blood + (tryptic soy agar with 0.5% arginine and 5% sheep blood).

### Library preparation and sequencing

Genomic DNA was prepared from anaerobic 24 hour liquid cultures using the media described above before extraction using the Powersoil DNA extraction kit (MoBio), PureLink genomic DNA minikit (ThermoFisher) or DNeasy UltraClean Microbial kit (Qiagen). Libraries were prepared with either an Apollo 324 instrument, the Nextera XT kit (Illumina), or TruSeq PCR-free kit (Illumina). Libraries were sequenced according to the platform and chemistry listed in **Supplemental Table 1**.

### Genome assembly and annotation

Demultiplexed sequences were analyzed first by FastQC before filter human and phiX174 contamination by removing reads that mapped to the hg19 and NC_001422.1 assemblies respectively with Bowtie 2 (40). Potential adapter contamination was removed using Trimmomatic (41) with additional sliding window filtering. Where paired reads were present, overlaps were assembled using VSEARCH (42) for use as a separate single-ended library for assembly. Assembly was carried out using the SPAdes genome assembler (43). Assembly statistics were generated using the QUAST quality assessment tool for genomes (44). Assembled genomes were submitted to NCBI for automated annotation using the Prokaryotic Genome Annotation Pipeline (PGAP). Assembled genomes were also annotated locally for use in comparative genomics analysis using Prokka (45). These annotated versions (as serialized general feature format files) have been packaged with ElenMatchR and can be extracted using the included unpackgenomes.R script. Phylogenetic analysis was carried out using PhyloPhlAn (46). Average nucleotide identity was calculated the Pyani package (widdowquinn.github.io/pyani/).

### Comparative genomic analysis

Clustering of gene orthologs were carried out using ProteinOrtho5 (32) across variable coverage and identity settings. For analysis in this manuscript, a minimal 60% identity and 80% coverage threshold were applied. To carry out gene-trait matching, gene presence was converted to a binary state, and used as the predictor matrix for the randomForest function of the random-Forest package in R. Results were ranked by mean decrease in GINI and plotted. The circular BLAST comparison was generated by splitting the *E. lenta* DSM 2243 type strain genome into successive 1000 bp ranges and aligning these against all strains (BLAST). These were then plotted in a circular representation colored by nucleotide identity. Genetic variation between DSM 2243 isolates was determined by using the tool Snippy (github.com/tseemann/snippy) and verified using manual inspection of read pileups and *breseq* (47). KEGG orthologous group content was assessed using GhostKOALA (48).

### Antibiotic resistance screening

Screening was carried out using Epsilometer (Etest) strips (bioMerieux) or broth dilution minimum inhibitory concentration (MIC) assay. For Etest strips, a 24 hour broth culture was diluted to an OD600 of 0.1 before 500 μL was spread on 135 mm plates with 4 strips per plate. The assays used were: CL 256, TC 256, KM 256, AM 256, VA 256, MX 32, PG 32, and MZ 256. Resistance was determined based on EuCAST breakpoint tables (version 8.0 2018) where available (31), or by MIC distribution. For the broth MIC assay, a 24h broth culture was inoculated at 1% v/v in a 96-well plate and incubated in anaerobic conditions at 37°C for 48h with OD600 recorded by an Eon microplate reader (BioTek). Strains were assayed twice and the mean value reported. Resistance was determined based on the bimodal distribution of MICs.

### Heterologous expression

The sequences of a putative tetW (DSM11767_02172) and aminoglycoside 3’-phosphotransferase (DSM11767_01344) were amplified from genomic DNA of *E. lenta* DSM 11767 with Q5 polymerase (NEB) using 500 μM primers (**Supplemental Table 7**) for 30 cycles. pET19btev (a derivative of pET19b with a tobacco etch virus [TEV] cut site replacing the enterokinase cut site) was used as the expression vector. Both inserts and plasmid were cut with NdeI and BamHI. The plasmid backbone was treated with recombinant shrimp alkaline phosphatase (rSAP) before both plasmid and inserts were gel purified (Qiaquick gel purification kit). Ligation was carried out with T4 ligase. All enzymes for cloning were purchased from New England Biolabs and used according to the manufacturer’s instructions. Ligation reactions were heat inactivated and transformed into *E. coli* DH5α and selected using LB media with 100μg/mL ampicillin. Plasmids were extracted from transformants and confirmed by sanger sequencing (GeneWiz) from the T7 promoter and terminator. Verified plasmids were then transformed into *E. coli* Rosetta and selected on LB with 50 μg/mL carbpenicillin (for pET19btev) and 30 μg/mL chloramphenicol (for selection of pRARE plasmid carrying rare codons). Verified transformants were then grown in LB broth for 8h with appropriate selection before being inoculated 0.5% v/v across a range of tetracycline, kanamycin, and IPTG concentrations in 96 well plates. Plates were incubated for 16 hours at 37°C and read at OD600 using an Eon microplate reader (BioTek).

### Mouse and human samples

Human samples were collected and analyzed as part of a diet-intervention study whose analysis is described elsewhere (clinical-trials.gov registration: NCT01105143). Human experiments were approved by the ethics committee of the Charité-Universitätsmedizin Berlin. Germ-free Swiss Webster mice were obtained from the UCSF Gnotobiotics core facility (gnotobiotics.ucsf.edu) and housed in gnotobiotic isolators (Class Biologically Clean). For the mono-colonization experiment, each of 10 Swiss Webster mice were orally gavaged with an inoculum of 1E8 CFUs of *E. lenta* DSM 2243 (prepared in an anaerobic environment and suspended in 200 μL PBS, which contained 0.05% L-cysteine-HCl). Mice were allowed to feed ad libitum on one of two isocaloric, isonitrogenous semi-purified diets composed of 10% sesame seeds (Teklad custom research diet). One diet was supplemented with 0.5% L-arginine (TD.150470) and provided to 5 mice; the other diet contained no supplemented L-arginine (TD.150471) and was provided to the other 5 mice. At 14 days following colonization, mice were euthanized, and the contents from each segment of the GI tract were collected. Arginine levels were not found to impact colonization (p>0.05, linear mixed effects model). For the 10-member mock community, 1E8 CFUs of each member (Supplemental Table 8) was combined and 1E8 CFUs (in 200 μL saline + 0.5% cysteine) was administered to 7 germ-free Swiss Webster mice who were given ad libitum accession to food (LabDiet 5021) and water. After 18 days, mice were sacrificed and contents collected from each segment of the GI tract. All mouse experiments were approved by the University of California San Francisco Institutional Animal Care and Use Committee. The mock community was profiled using 515F/806R golay-barcodes (49) and denoised using DADA2 (50).

### DNA extraction and qPCR

DNA was extracted from 100 mg aliquots of material using the ZymoBIOMICS 96 MagBead DNA Kit (Zymo D4302) according to the manufacturer’s protocol with an additional 10 minute incubation after mechanical cell disruption at 65°C. Disruption was carried out using a BioSpec Mini-Beadbeater-96 for 5 minutes. qPCR was carried out using Taq Universal Probes Supermix (BioRad 1725131), in a CFX384 thermocycler (BioRad) in triplicate 10 μL reactions with all oligos (Supplemental Table 7) present at 200 nM final concentration and 4 μL of purified gDNA. DNA was amplified using 40 cycles of 5 seconds at 95°C and 30 seconds at 60°C after 5 minutes initial denaturation at 95°C. Absolute copies were determined by comparison against a standard curve of *E. lenta* DSM2243 genomic DNA. The primer efficiencies of the *elnmrk1* and *cgr2* assays are 105% and 103.5% respectively. A practical limit of quantification of 1400 genome equivalents/ml was determined as it was the last dilution in the series fitting the linear trend-line with concordant detection in all replicate wells.

## Supplemental Figures

**Supplemental Figure 1.**
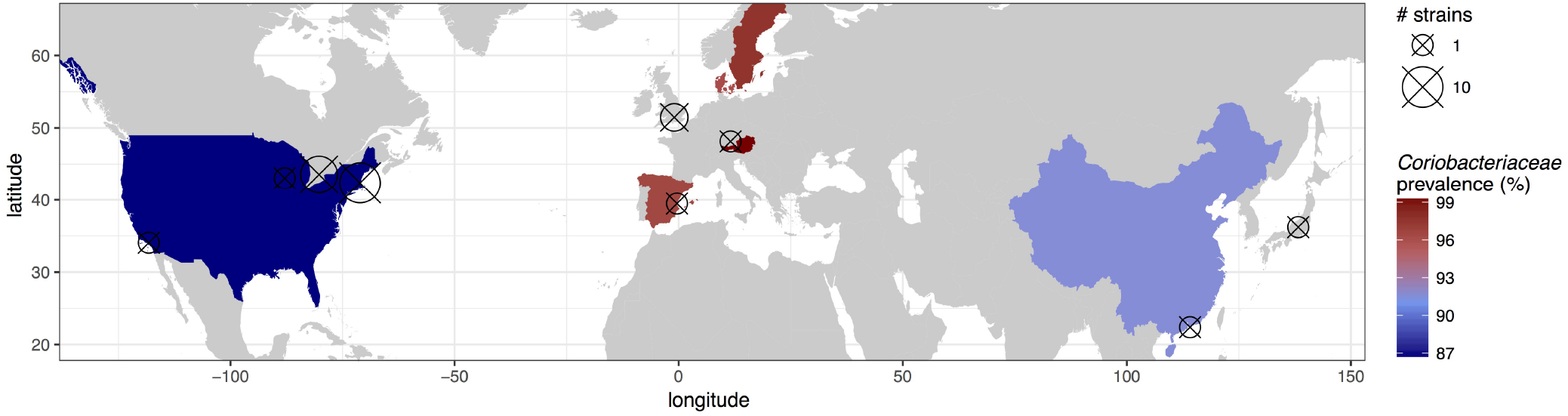
Isolation locations and Coriobacteriia prevalence. Location of isolation where known is plotted for the strain collection and prevalence of Coriobacteriia (as determined from MetaPhlAn2 analysis of curated shotgun metagenomes (36)) is plotted where known. Note: *Coriobacteriaceae* prevalence estimates were used as the MetaPhlAn2 database is based on the previously accepted nomenclature where in all members of the Coriobacteriia were within the family *Coriobacteriaceae*.

**Supplemental Figure 2.**
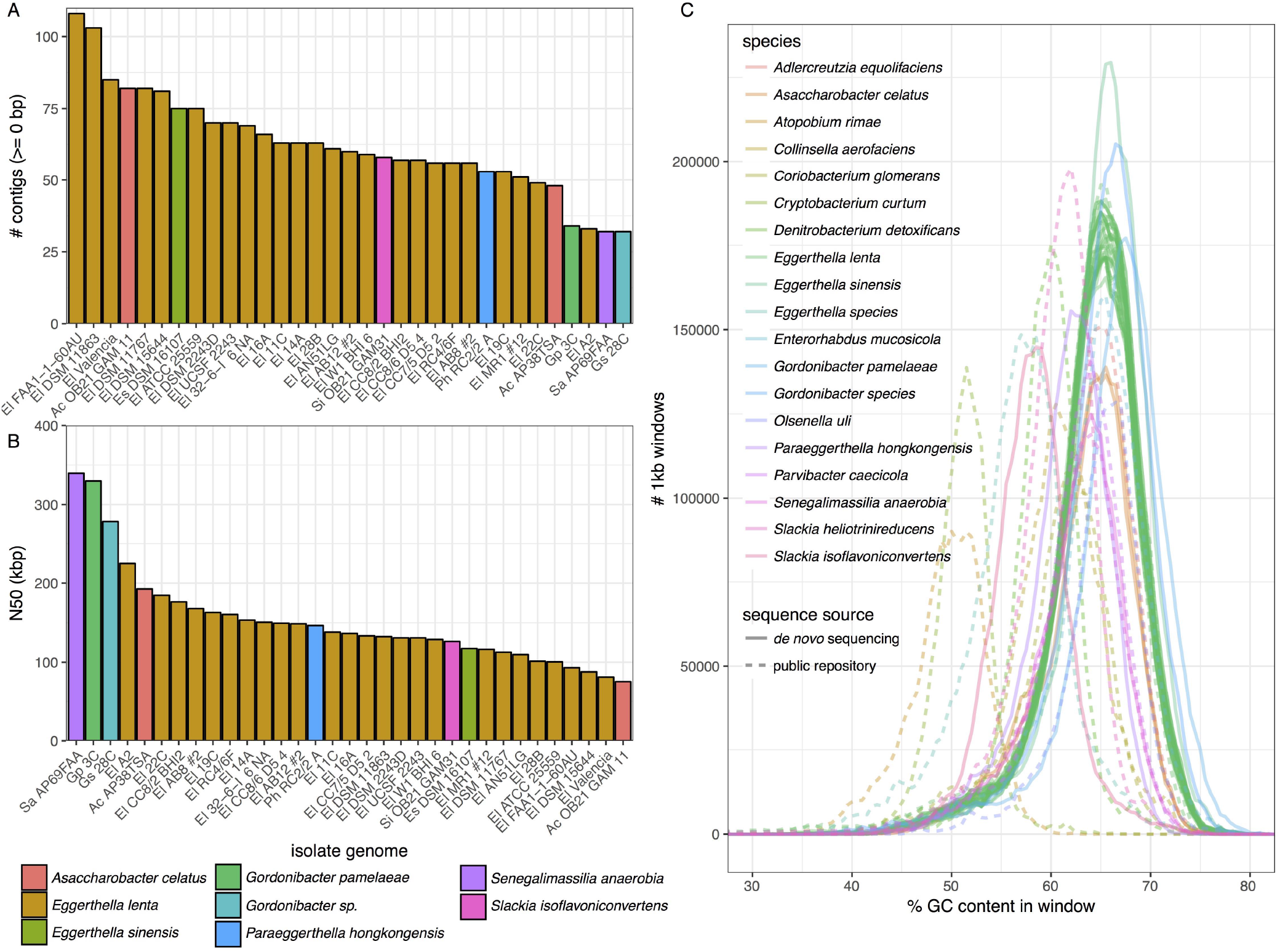
Sequence assembly statistics. High quality draft genomes were generated for newly sequenced strains based on: (**A**) a small number of contigs per genome (62 ±18 mean ± sd), and (**B**) a large contig N50 (length of contig of which >50% genome length is contained in, 152 ± 62 kbp). (**C**) GC content distribution by genome demonstrates and approximately Gaussian distribution within any given genome indicating genome purity.

**Supplemental Figure 3.**
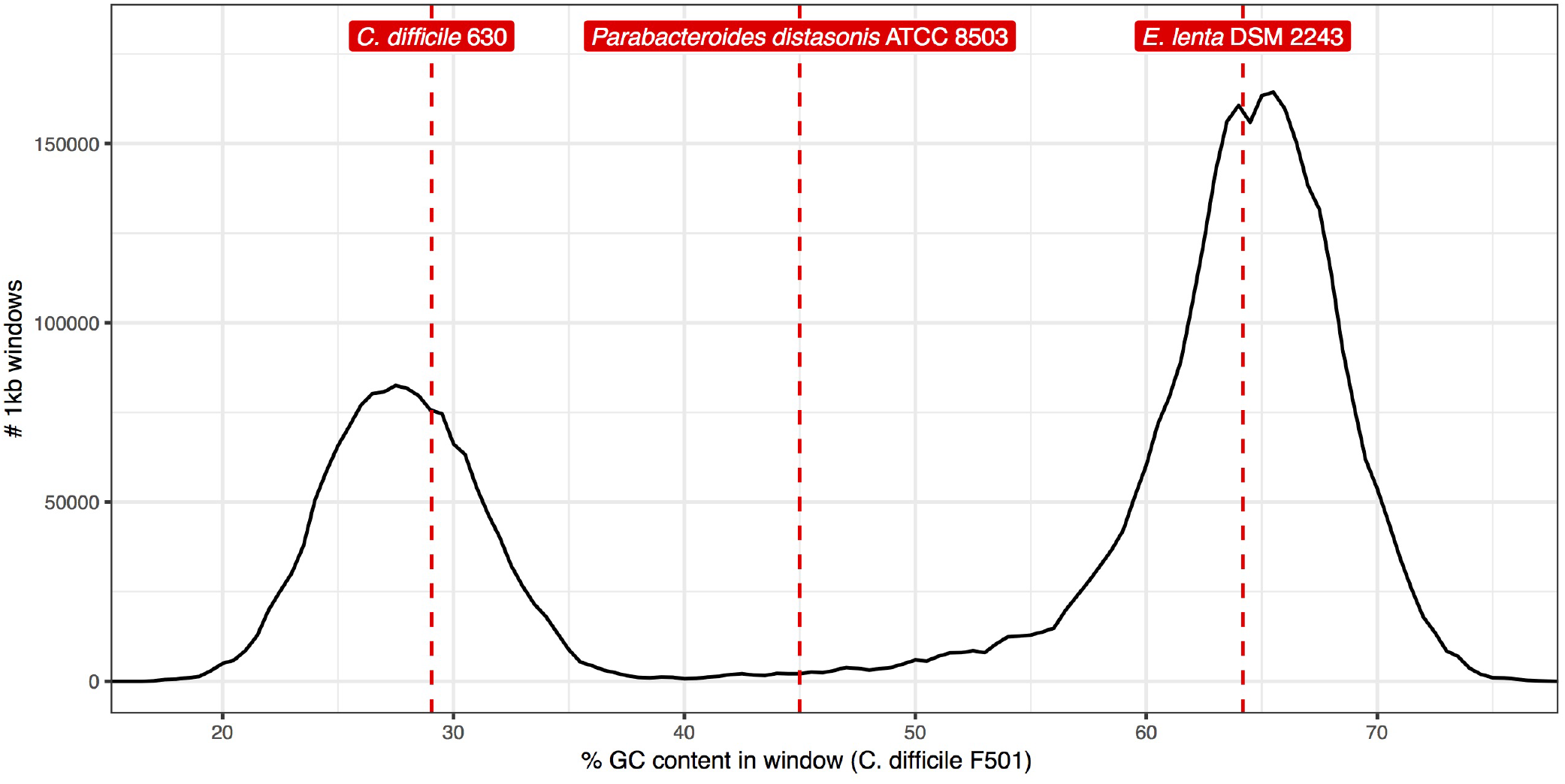
Identification of genomic DNA contamination in the *C. difficile* F501 genome. Rather than a Gaussian distribution of GC content (examples in **Supplemental Figure 2**), GC content is bimodal and consistent with a high degree of *E. lenta* in sequenced genomic DNA preparation. Recovery of 16S rRNA sequences from contigs suggests the presence of both *E. lenta* and *P. distasonis* in the genome (their average GC contents are denoted by dashed red lines).

**Supplemental Figure 4.**
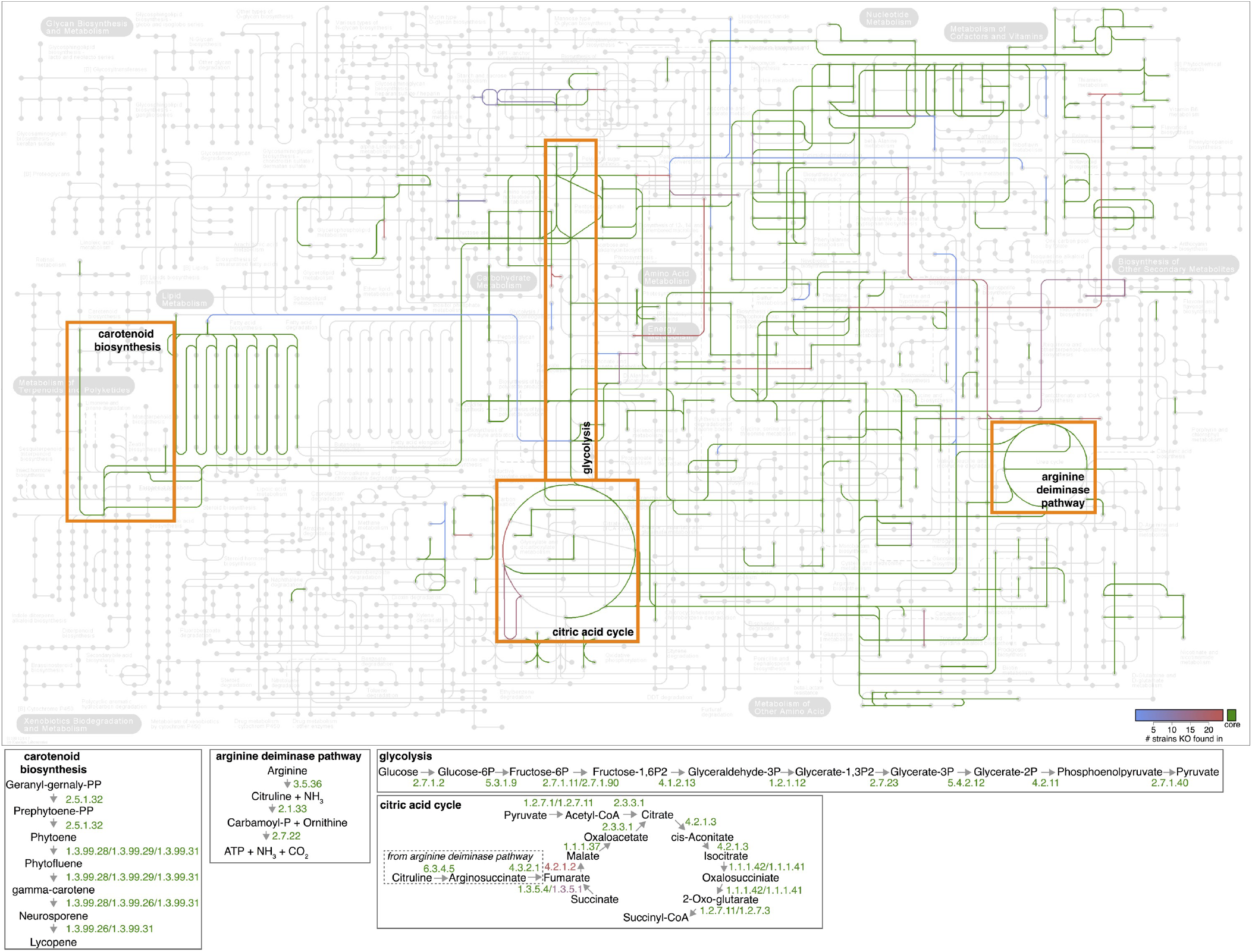
Conservation of core metabolic pathways. Conservation of KEGG orthologies (KOs) is plotted in the context of global metabolism (map accession: ko01100) with selected pathways displayed below with intermediates and enzymes (by enzyme commission number).

**Supplemental Figure 5.**
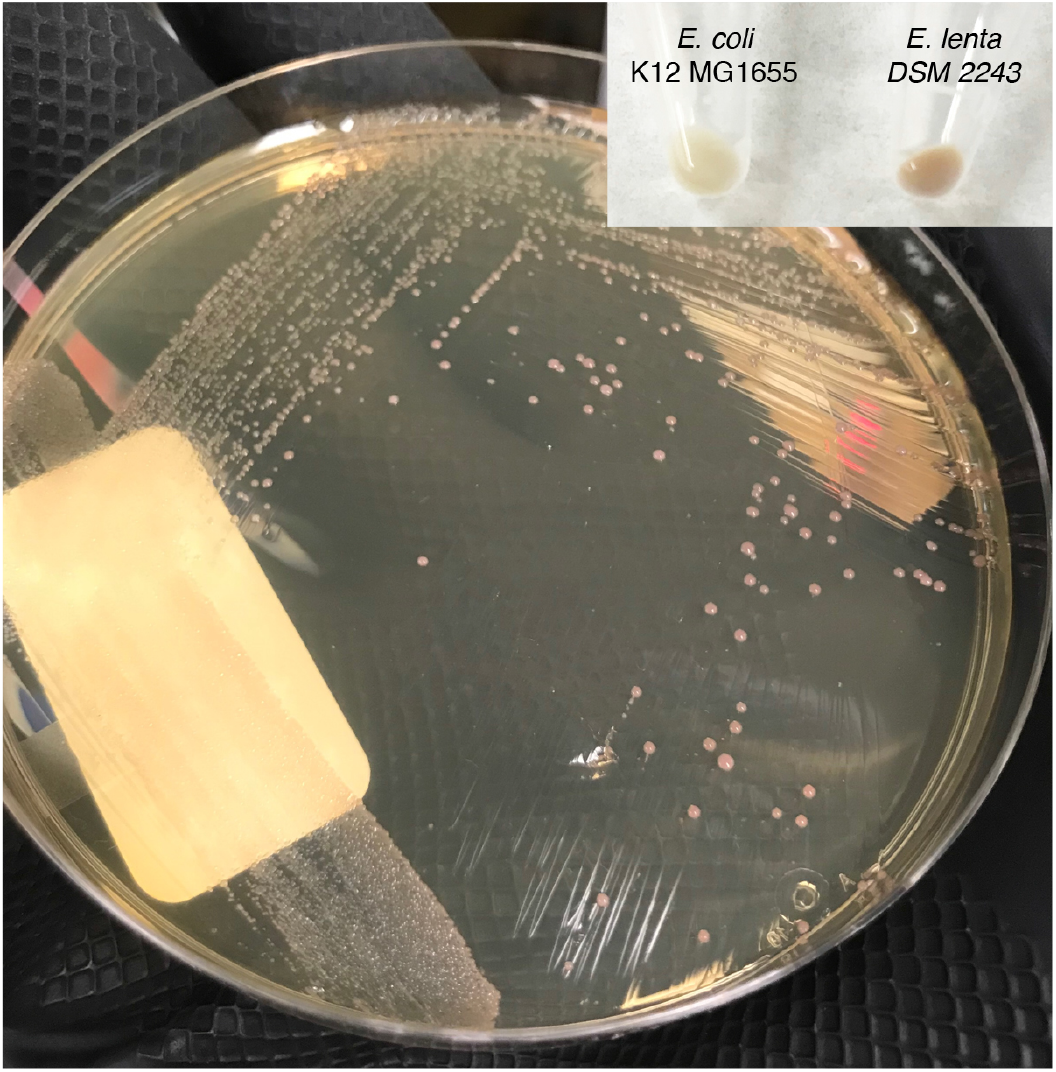
Pigmentation in *E. lenta* DSM 2243. This culture was streaked from frozen glycerol stock on to BHI++ (see methods) and incubated for 72h in anaerobic conditions resulting in a typical red-brown pigment production (inset pellet from 24h anaerobic broth culture in BHI +1% arginine with *E. coli* as comparison).

**Supplemental Figure 6.**
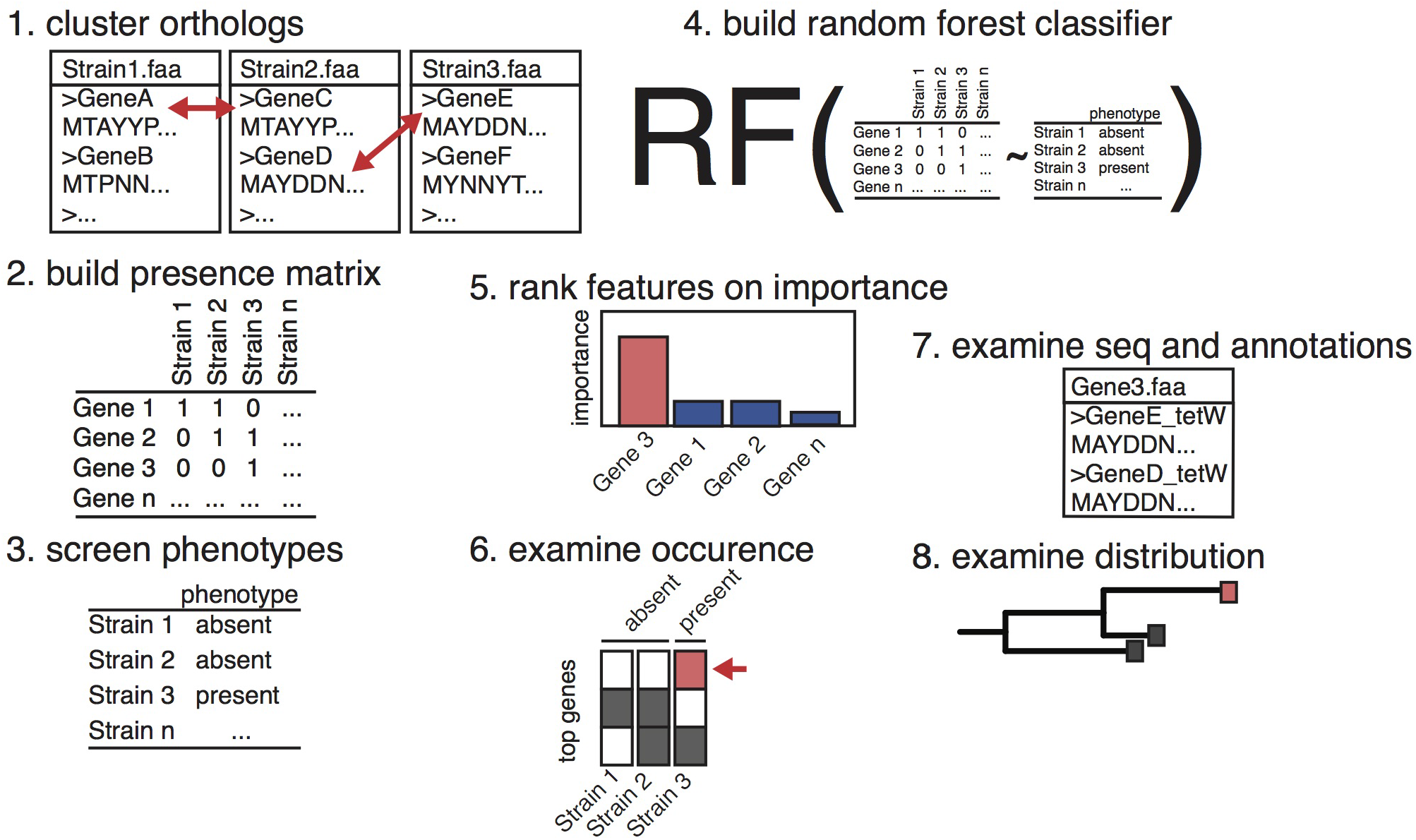
ElenMatchR algorithm. Precomputed matrices of gene presence/abundance are binarized and used as the input for a random forest classifier. The trained model is then used to rank features on importance extracting the top features for user interpretation of biological significance by heat maps of presence/absence, annotation tables, raw sequence for downstream analysis, phylogenetic and pan-genome trees.

**Supplemental Figure 7.**
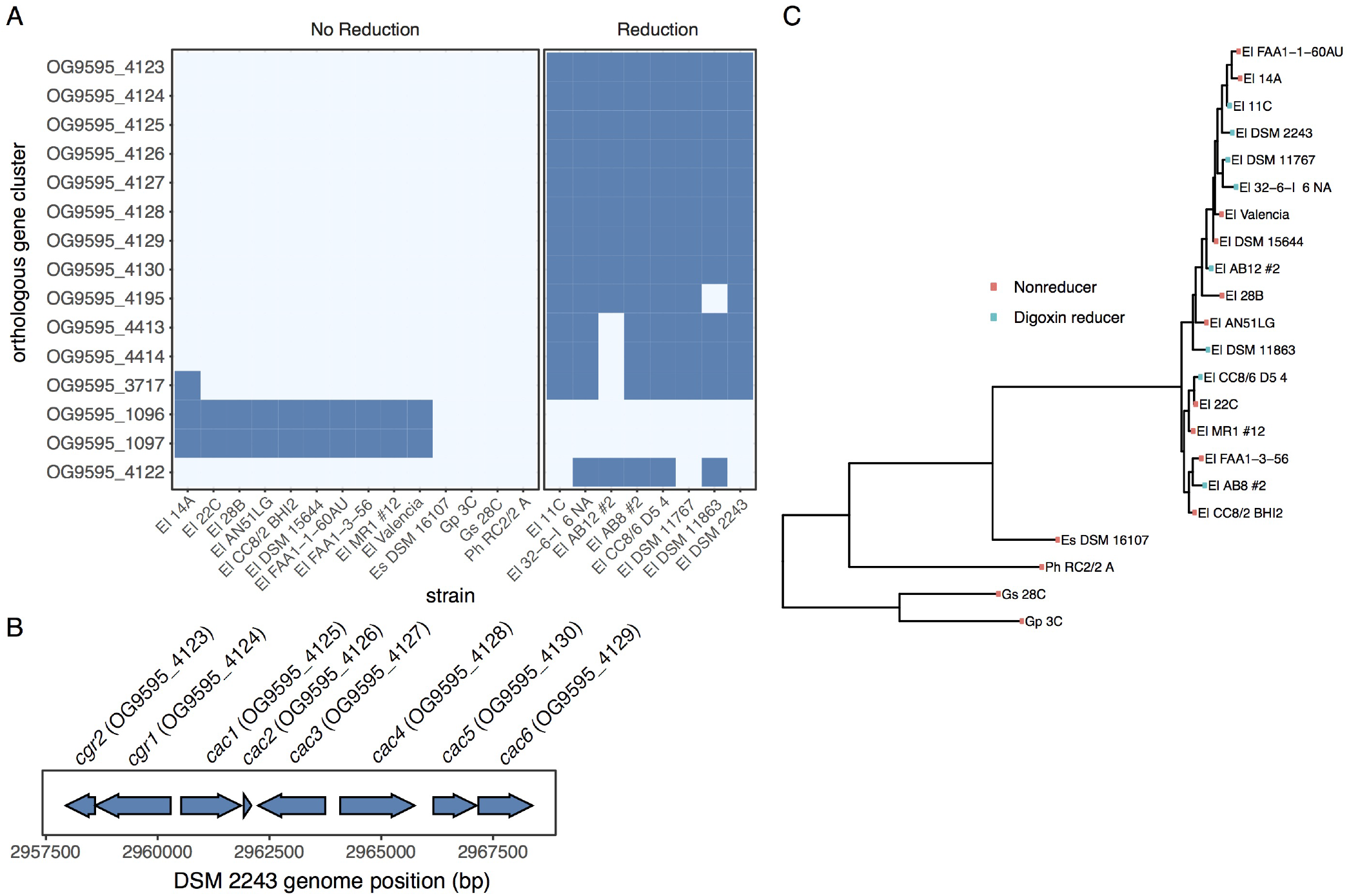
Identification of cardiac glycoside reductase operon by ElenMatchR. (**A**) Analysis of the occurrence of the top 15 predictive features for digoxin metabolism reveals 8 genes which perfectly co-occur with the ability to reduce digoxin. (**B**) Using supplied annotation data, it is revealed that the 8 genes occur in a single locus termed which we have term the cgr-associated gene cluster. (**C**) Occurence of the digoxin reduction phenotype in a phylogenetic tree demonstrates a lack of phylogenetic signal.

## References

1. Human Microbiome Project Consortium. A catalog of reference genomes from the human microbiome. Science. 2010;328(5981):994–999.

2. Wu D, Hugenholtz P, Mavromatis K, Pukall R, Dalin E, Ivanova NN, et al. A phylogeny-driven genomic encyclopaedia of Bacteria and Archaea. Nature. 2009;462(7276):1056.

3. Wayne LG, Brenner DJ, Colwell RR, Grimont PA, Kandler O, Krichevsky MI, et al. Report of the ad hoc committee on reconciliation of approaches to bacterial systematics. International Journal of Systematic Bacteriology. 1987;37(4):463–464.

4. Kim M, Oh HS, Park SC, Chun J. Towards a taxonomic coherence between average nucleotide identity and 16S rRNA gene sequence similarity for species demarcation of prokaryotes. International Journal of Systematic and Evolutionary Microbiology. 2014;64(2):346–351.

5. Quintero AC, Konstantinidis KT. Bacterial species may exist, metagenomics reveal. Environmental Microbiology. 2012;14(2):347–355.

6. Sybesma W, Molenaar D, van IJcken W, Venema K, Kort R. Genome instability in *Lactobacillus rhamnosus* GG. Applied and Environmental Microbiology. 2013;79(7):2233–2239.

7. Langille MGI, Zaneveld J, Caporaso JG, McDonald D, Knights D, Reyes JA, et al. Predictive functional profiling of microbial communities using 16S rRNA marker gene sequences. Nature Biotechnology. 2013;31(9):814–821.

8. Haas F, König H. *Coriobacterium glomerans* gen. nov., sp. nov. from the Intestinal Tract of the Red Soldier Bug. International Journal of Systematic and Evolutionary Microbiology. 1988;38(4):382–384.

9. Rosenberg E, DeLong EF, Lory S, Stackebrandt E, Thompson F, editors. The Prokaryotes. 4th ed. Actinobacteria. Berlin, Heidelberg: Springer Berlin Heidelberg; 2014.

10. Thota VR, Dacha S, Natarajan A, Nerad J. *Eggerthella lenta* bacteremia in a Crohn’s disease patient after ileocecal resection. Future Microbiology. 2011;6(5):595–597.

11. Gardiner BJ, Tai AY, Kotsanas D, Francis MJ, Roberts SA, Ballard SA, et al. Clinical and Microbiological Characteristics of *Eggerthella lenta* Bacteremia. Journal of Clinical Microbiology. 2015;53(2):626–635.

12. Venugopal AA, Szpunar S, Johnson LB. Risk and prognostic factors among patients with bacteremia due to *Eggerthella lenta*. Anaerobe. 2012;18(4):475–478.

13. Qin J, Li Y, Cai Z, Li S, Zhu J, Zhang F, et al. A metagenome-wide association study of gut microbiota in type 2 diabetes. Nature. 2012;490(7418):55.

14. Jie Z, Xia H, Zhong SL, Feng Q, Li S, Liang S, et al. The gut microbiome in atherosclerotic cardiovascular disease. Nature Communications. 2017;8(1):845.

15. Chen J, Wright K, Davis JM, Jeraldo P, Marietta EV, Murray J, et al. An expansion of rare lineage intestinal microbes characterizes rheumatoid arthritis. Genome Medicine. 2016;8(1):43.

16. Haiser HJ, Gootenberg DB, Chatman K, Sirasani G, Balskus EP, Turnbaugh PJ. Predicting and manipulating cardiac drug inactivation by the human gut bacterium *Eggerthella lenta*. Science. 2013;341(6143):295–298.

17. Haiser HJ, Seim KL, Balskus EP, Turnbaugh PJ. Mechanistic insight into digoxin inactivation by *Eggerthella lenta* augments our understanding of its pharmacokinetics. Gut Microbes. 2014;5(2):233–238.

18. Koppel N, Bisanz JE, Pandelia ME, Turnbaugh PJ, Balskus EP. Discovery and characterization of a prevalent human gut bacterial enzyme sufficient for the inactivation of plant toxins. eLife. 2018;In Press.

19. Devlin AS, Fischbach MA. A biosynthetic pathway for a prominent class of microbiota-derived bile acids. Nature Chemical Biology. 2015;11(9):685–690.

20. Harris SC, Devendran S, Méndez-García C, Mythen SM, Wright CL, Fields CJ, et al. Bile acid oxidation by *Eggerthella lenta* strains C592 and DSM 2243T. Gut Microbes. 2018;In Press.

21. Clavel T, Doré J, Blaut M. Bioavailability of lignans in human subjects. Nutrition Research Reviews. 2006;19(2):187–196.

22. Yokoyama Si, Oshima K, Nomura I, Hattori M, Suzuki T. Complete genomic sequence of the equol-producing bacterium*Eggerthella* sp. strain YY7918, isolated from adult human intestine. Journal of Bacteriology. 2011;193(19):5570–5571.

23. Matthies A, Loh G, Blaut M, Braune A. Daidzein and Genistein Are Converted to Equol and 5-Hydroxy-Equol by Human Intestinal *Slackia isoflavoniconvertens* in Gnotobiotic Rats. The Journal of Nutrition. 2012;142(1):40–46.

24. König H. Coriobacteriia class. nov. In Bergey’s Manual of Systematics of Archaea and Bacteria. 2015; p. 10.1002–9781118960608.cbm00005.

25. Kumar K, Jaiswal SK, Dhoke GV, Srivastava GN, Sharma AK, Sharma VK. Mechanistic and structural insight into promiscuity based metabolism of cardiac drug digoxin by gut microbial enzyme. Journal of Cellular Biochemistry. 2018;https://doi.org/10.1002/jcb.26638.

26. Parte AC. LPSN—list of prokaryotic names with standing in nomenclature. Nucleic Acids Research. 2014;42(D1):D613–D616.

27. Truong DT, Franzosa EA, Tickle TL, Scholz M, Weingart G, Pasolli E, et al. MetaPhlAn2 for enhanced metagenomic taxonomic profiling. Nature. 2015;12(10):902–903.

28. Minamida K, Ota K, Nishimukai M, Tanaka M, Abe A, Sone T, et al. *Asaccharobacter celatus* gen. nov., sp. nov., isolated from rat caecum. International Journal of Systematic and Evolutionary Microbiology. 2008;58(5):1238–1240.

29. Maruo T, Sakamoto M, Ito C, Toda T, Benno Y.*Adlercreutzia equolifaciens* gen. nov., sp. nov., an equol-producing bacterium isolated from human faeces, and emended description of the genus*Eggerthella*. International Journal of Systematic and Evolutionary Microbiology. 2008;58(5):1221–1227.

30. Saunders E, Pukall R, Abt B, Lapidus A, Del Rio TG, Copeland A, et al. Complete genome sequence of *Eggerthella lenta* type strain (VPI 0255 T). Standards in Genomic Sciences. 2009;1(2):174–182.

31. European Committee on Antimicrobial Susceptibility Testing. Breakpoint tables for interpretation of MICs and zone diameters. version 80. 2018; p. gram–positive anaerobes.

32. Lechner M, Findeiß S, Steiner L, Marz M, Stadler PF, Prohaska SJ. Proteinortho: Detection of (Co-)orthologs in large-scale analysis. BMC Bioinformatics. 2011;12(1):124.

33. Nupur LNU, Vats A, Dhanda SK, Raghava GPS, Pinnaka AK, Kumar A. ProCarDB: a database of bacterial carotenoids. BMC Microbiology. 2016;16(1):96.

34. Juhas M, Crook DW, Hood DW. Type IV secretion systems: tools of bacterial horizontal gene transfer and virulence. Cellular Microbiology. 2008;10(12):2377–2386.

35. Song L, Pan Y, Chen S, Zhang X. Structural characteristics of genomic islands associated with GMP synthases as integration hotspot among sequenced microbial genomes. Computational Biology and Chemistry. 2012;36:62–70.

36. Nayfach S, Fischbach MA, Pollard KS. MetaQuery: a web server for rapid annotation and quantitative analysis of specific genes in the human gut microbiome. Bioinformatics. 2015;31(20):3368–3370.

37. Sunagawa S, Mende DR, Zeller G, Izquierdo-Carrasco F, Berger SA, Kultima JR, et al. Metagenomic species profiling using universal phylogenetic marker genes. Nature Methods. 2013;10(12):1196–1199.

38. Bayjanov JR, Molenaar D, Tzeneva V, Siezen RJ, van Hijum SA. PhenoLink - a web-tool for linking phenotype to ~ omics data for bacteria: application to gene-trait matching for *Lactobacillus plantarum* strains. BMC Genomics. 2012;13(1):170.

39. McArthur AG, Waglechner N, Nizam F, Yan A, Azad MA, Baylay AJ, et al. The comprehensive antibiotic resistance database. Antimicrobial Agents and Chemotherapy. 2013;57(7):3348–3357.

40. Langmead B, Salzberg SL. Fast gapped-read alignment with Bowtie 2. Nature Methods. 2012;9(4):357–359.

41. Bolger AM, Lohse M, Usadel B. Trimmomatic: a flexible trimmer for Illumina sequence data. Bioinformatics. 2014;30(15):2114–2120.

42. Rognes T, Flouri T, Nichols B, Quince C, Mahé F. VSEARCH: a versatile open source tool for metagenomics. PeerJ. 2016;4:e2584.

43. Bankevich A, Nurk S, Antipov D, Gurevich AA, Dvorkin M, Kulikov AS, et al. SPAdes: A New Genome Assembly Algorithm and Its Applications to Single-Cell Sequencing. Journal of Computational Biology. 2012;19(5):455–477.

44. Gurevich A, Saveliev V, Vyahhi N, Tesler G. QUAST: quality assessment tool for genome assemblies. Bioinformatics. 2013;29(8):1072–1075.

45. Seemann T. Prokka: rapid prokaryotic genome annotation. Bioinformatics. 2014;30(14):2068–2069.

46. Segata N, Börnigen D, Morgan XC, Huttenhower C. PhyloPhlAn is a new method for improved phylogenetic and taxonomic placement of microbes. Nature Communications. 2013;4:2304.

47. Deatherage DE, Barrick JE. Identification of mutations in laboratory-evolved microbes from next-generation sequencing data using breseq. Methods in Molecular Biology. 2014;1151(Chapter 12):165–188.

48. Kanehisa M, Sato Y, Morishima K. BlastKOALA and GhostKOALA: KEGG Tools for Functional Characterization of Genome and Metagenome Sequences. Journal of Molecular Biology. 2016;428(4):726–731.

49. Caporaso JG, Lauber CL, Walters WA, Berg-Lyons D, Huntley J, Fierer N, et al. Ultra-high-throughput microbial community analysis on the Illumina HiSeq and MiSeq platforms. The ISME Journal. 2012;6(8):1621–1624.

50. Callahan BJ, McMurdie PJ, Rosen MJ, Han AW, Johnson AJA, Holmes SP. DADA2: High-resolution sample inference from Illumina amplicon data. Nature Methods. 2016;13(7):581–583.

